# Comprehensive Proteomic Quantification of Bladder Stone Progression in a Cystinuric Mouse Model Using Data-Independent Acquisitions

**DOI:** 10.1101/2021.04.06.438573

**Authors:** Jacob Rose, Nathan Basisty, Tiffany Zee, Cameron Wehrfritz, Neelanjan Bose, Pierre-Yves Desprez, Pankaj Kapahi, Marshall Stoller, Birgit Schilling

## Abstract

Cystinuria is one of various disorders that cause biomineralization in the urinary system, including bladder stone formation in humans. It is most prevalent in children and adolescents and more aggressive in males. There is no cure, and only limited disease management techniques help to solubilize the stones. Recurrence, even after treatment, occurs frequently. Other than a buildup of cystine, little is known about factors involved in the formation, expansion, and recurrence of these stones. This study sought to define the growth of bladder stones, guided by micro-computed tomography imaging, and to profile dynamic stone proteome changes in a cystinuria mouse model. After bladder stones developed *in vivo*, they were harvested and separated into four developmental stages (sand, small, medium and large stone), based on their size. Data-dependent and data-independent acquisitions allowed deep profiling of stone proteomics. The proteomic signatures and pathways illustrated major changes as the stones grew. Stones initiate from a small nidus, grow outward, and show major enrichment in ribosomal proteins and factors related to coagulation and platelet degranulation, suggesting a major dysregulation in specific pathways that can be targeted for new therapeutic options.

## Introduction

Cystinuria is a genetic disorder characterized by aggressive/recurrent kidney stone formation. It is caused by mutations in the solute carrier family 3 member 1 (*SLC3A1*), solute carrier family 7 member 9 (*SLC7A9*), and/or in the recently identified *AGT1* gene that codes for a cystine reabsorption transporter [1–3]. Patients with cystinuria typically excrete markedly elevated levels of quantitative urinary cystine and develop cystinuric stones at a high rate of recurrence due to the low solubility of cystine. Current interventions aim to decrease urinary cystine concentration with a combination of the following: i) increased fluid intake, ii) a low protein diet, and iii) urine alkalinizing drugs or cystine-binding thiol drugs, such as penicillamine and tiopronin [4]. In spite of these interventions, cystinuric patients experience high stone recurrence rates and endure repeated surgical interventions. Furthermore, these medications can be associated with serious adverse side effects, including decreased renal function [5].

Little is known about the factors that contribute to the severity and recurrence rates of cystinuria-related stone events [6–8]. While some cystinuric patients experience chronic stone formation with multiple surgeries per year, others have few or no stone events throughout their lifetime; this is frequently independent of their quantitative urinary cystine levels. Even when cystinuric patients present above a urine threshold of 300 mg/L cystine output/day, there is typically no correlation between cystine excretion and recurrence rate of stone formation [9]. Cystine output clinically correlates with the ability to reabsorb cystines, and increased cystine output is a risk factor for stone development. Pharmaceutical interventions that target urinary cystine output have had limited efficacy on stone recurrence. These observations suggest that more factors contribute to cystine stone development.

There are only few reports on urinary stone proteomics, but the human urine proteome is well characterized, and large repositories are provided by the Human Kidney and Urine Proteome Project (http://www.hkupp.org/). Human urine proteomics is a promising approach for biomarker discovery, particularly for studying the pathogenesis of kidney diseases and diseases of the urothelial tract [10]. Additionally, proteomic studies of mouse urine show similar protein signatures when age-matched with humans [11]. Cystinuric models are not as well understood, and proteomic studies for patients with cystinuria are lacking. Some studies have been conducted, mostly in children [12]; however, they examined urine rather than the corresponding stones, leaving the stone protein compositions unknown [13].

Urinary proteins are generally believed to contribute to the development of urinary stones by promoting crystal aggregation and adherence to the renal epithelium [14]. Matrix proteins have been detected in proteomic profiles of human urinary stones, but due to radiation concerns and poor resolution with clinically available CT imaging, especially in comparison to micro-CT utilized in rodent animal models, scarce information is available on the early events that lead to urinary stone development and the progression of kidney stones. Thus, analysis has been limited to larger, mature stones that have been surgically extracted from patients presenting with acute renal colic.

Using a cystinuric mouse model, a dynamic study on cystinuric stone formation beginning with the smallest aggregates or “sand” to the largest stones categorizing into three other separate sizes was undertaken. For accurate quantification of relative protein abundance, we used label-free proteomic data-independent acquisitions (DIA) or SWATH assays [15, 16] that allow us to accurately determine changes in relative protein expression levels between multiple different sample sets, specifically comparing protein profiles between the different stone sizes and categories. This comprehensive DIA technology provides high sensitivity to quantify changes in relative protein abundance during stone formation. This report highlights the complex organic composition of bladder stones with a constantly changing proteome as stones increase in size. We provide molecular insight into the initiation and growth of these stones, as well as valuable comparisons to human urine composition and kidney disorders.

## Materials and Methods

### Reagents and standards

HPLC solvents (e.g., acetonitrile and water) were obtained from Burdick & Jackson (Muskegon, MI). Reagents for protein chemistry (e.g., iodoacetamide, dithiothreitol (DTT), ammonium bicarbonate, formic acid, and urea) were purchased from Sigma Aldrich (St. Louis, MO). Proteomics grade trypsin was from Promega (Madison WI). HLB Oasis SPE cartridges were purchased from Waters (Milford, MA).

### Mice and micro-computed tomography

All procedures and protocols were approved by the Institutional Animal Care and Use Committee of the Buck Institute for Research on Aging. Male *Slc3a1^-/-^* mice (6–12 weeks old) were anesthetized with isoflurane and scanned using Skyscan 1176 μCT scanner (Bruker Corp, Billerica, MA). The Skyscan reconstruction program NRecon was used for image reconstruction, and bladder stone volume was quantified using the Bruker CT-Analyzer (CTAn, Version 1.14) program with Hounsfield units. 3-D image models were created using CTAn and Bruker CT-Volume (CTVol, Version 2.2).

### Stone collection and sample preparation

Batches of cystine sand and stone samples were harvested from the bladder of male *Slc3a1^-/-^* mice in at least four biological replicates. Sand and stone samples were categorized, according to their aggregate surface characteristics and apparent diameters. Sand samples were granular and small in diameter (<1 mm^2^), and stone samples were lithic and larger in diameter (>5 mm^2^). All sand and stone samples were washed thoroughly in deionized water to remove contaminants and debris. Dried samples were then ground into a powder in a mortar and pestle. To extract proteins from the sand and stone samples, we used a modified version of the protocol of Jiang *et al.* [17]. Briefly, ground stone and sand samples were lysed in 6 M guanidine HCl, 100 mM Tris (pH 7.4), and Sigma-Aldrich complete EDTA-free protease inhibitor cocktail. The samples were incubated at 4°C for 72 hours, and the supernatant was collected.

### Cystine determination of stones

Cystine sand and stone samples that were harvested from the bladder of male *Slc3a1^-/-^* mice were washed thoroughly in deionized water. Dry samples were then ground into a powder in a mortar and pestle. Samples were sonicated and solubilized in water at 37°C for 16 hours with shaking, and the supernatant was collected for analysis.

### Sample processing for mass spectrometry

The protein mixture (typically 100 μg protein lysate) was reduced with 20 mM DTT (37°C for 1 hour), and then alkylated with 40 mM iodoacetamide (30 min at RT in the dark). Samples were diluted 10-fold with 100 mM Tris, pH 8.0, and incubated overnight at 37°C with sequencing grade trypsin (Promega) added at a 1:50 enzyme:substrate ratio (wt/wt). Samples were then acidified with formic acid and desalted using HLB Oasis SPE cartridges (Waters). Proteolytic peptides were eluted, concentrated to near dryness by vacuum centrifugation, re-suspended and further desalted (C-18 zip-tips) for insoluble protein mass spectrometric analysis.

### Mass spectrometry acquisition and analysis

Samples were analyzed by reverse-phase HPLC-ESI-MS/MS with an Eksigent Ultra Plus nano-LC 2D HPLC system (Dublin, CA), combined with a cHiPLC System, and directly connected to a quadrupole time-of-flight TripleTOF 6600 (QqTOF) mass spectrometer (SCIEX). Briefly, DIA acquisitions acquire MS/MS fragment ions from essentially all peptide precursor ions by consecutively passing large *m/z* DIA segments over the MS1 scan range and acquiring high-resolution MS/MS scans for each Δm/z segment. Typically, we scan from *m/z* 400-1250 with 64 variable window m/z segments [18–20]. Typically, mass resolution for MS1 scans and corresponding precursor ions was ~45,000, and resolution for MS/MS scans and resulting fragment ions was ~15,000 (‘high-sensitivity’ product ion scan mode). For acquisition, the autosampler was operated in full injection mode overfilling a 3-μL loop with 4 μL of analyte for optimal sample delivery reproducibility. Briefly, after injection, peptide mixtures were transferred onto a trap chip (with 200 μm x 6 mm ChromXP C18-CL chip, 3 μm, 300 Å, SCIEX) and washed at 2 μL/min for 10 min with the loading solvent (H_2_O/0.1% formic acid). Subsequently, peptides were transferred to each 75 μm x 15 cm ChromXP C18-CL chip, 3 μm, 300 Å, (SCIEX), and eluted at a flow rate of 300 nL/min with the following gradient: at 5% solvent B in A (from 0–5 min), 5–8% solvent B in A (from 5–12 min), 8–35% solvent B in A (from 12–67 min), 35–80% solvent B in A (from 67–77 min), at 80% solvent B in A (from 77–87 min), with a total runtime of 120 min, including mobile phase equilibration. Solvents were prepared as follows, mobile phase A: 2% acetonitrile/98% of 0.1% formic acid (v/v) in water, and mobile phase B: 98% acetonitrile/2% of 0.1% formic acid (v/v) in water.

Data-dependent acquisitions (DDA) were performed on the TripleTOF 6600 to obtain MS/MS spectra for the 30 most abundant precursor ions (100 msec per MS/MS) after each survey MS1 scan (250 msec), yielding a total cycle time of 3.3 sec as described [21, 22]. For collision-induced dissociation tandem mass spectrometry (CID-MS/MS), the mass window for precursor ion selection of the quadrupole mass analyzer was set to ± 1 m/z. Each of the 21 stone samples was acquired in two technical replicates (MS injection replicates) using the Analyst 1.7 (build 96) software. All database search results and details for peptide identifications are provided in **S3 Table.** For additional quantitative assessments data-independent acquisitions (DIA), or SWATH acquisitions, were performed for four ‘sand’ stone samples and four large stone samples, plus additional small (2x) and medium (2x) stone samples. Briefly, instead of the Q1 quadrupole transmitting a narrow mass range through to the collision cell, a wider window of variable window width (5–90 *m/z*) is passed in incremental steps over the full mass range (*m/z* 400–1250 with 64 DIA segments, 45 msec accumulation time each, yielding a cycle time of 3.2 sec that includes one MS1 scan with 250 msec accumulation time). The variable window width is adjusted, according to the complexity of the typical MS1 ion current observed within a certain *m/z* range using a SCIEX ‘variable window calculator’ algorithm (more narrow windows were chosen in ‘busy’ *m/z* ranges, wide windows in *m/z* ranges with few eluting precursor ions). DIA workflows produce complex MS/MS spectra, which are a composite of all the analytes within each selected Q1 *m/z* window.

### Mass spectrometry database search

Mass spectrometric data was searched using the database search engine Protein Pilot [23] (SCIEX 5.0, revision 4769) with the Paragon algorithm (5.0.0.0.4767). The search parameters were set as follows: trypsin digestion, cysteine alkylation set to iodoacetamide, and *Mus musculus* as species (33,338 protein entries, SwissProt database release 2014_05). Additional searches utilized phosphorylation emphasis. Trypsin specificity was assumed as C-terminal cleavage at lysine and arginine. Processing parameters were set to “Biological modification” and a thorough ID search effort was used. For database searches, a cut-off peptide confidence value of 99 was chosen, and a minimum of two identified peptides per protein was required. The Protein Pilot False Discovery Rate (FDR) analysis tool, and the Proteomics System Performance Evaluation Pipeline (PSPEP) algorithm provided a global FDR of 1% and a local FDR at 1% in all cases.

### Mass spectrometry data processing and quantification

#### Quantitative processing of DIA data

DIA data was processed with Spectronaut Pulsar v12 (12.020491.3.1543) software [24] from Biognosys, using a spectral ion library generated from the data-dependent acquisitions. In addition, we used Skyline 3.5 [25], an open source software project (http://proteome.gs.washington.edu/software/skyline), to process DIA data for relative quantitation comparing large stone and sand samples. For the DIA MS2 data sets, Skyline quantitation was based on XICs of up to 10 MS/MS fragment ions, typically y- and b-ions, matching to specific peptides in the spectral libraries used. Significance was assessed using q-values from two-tailed t-tests adjusted for multiple testing corrections. Significantly changed proteins were accepted at a 5% FDR (q-value < 0.05). All database search results and details for peptide identifications are provided in **S1 Table.** Pathway enrichments were determined with the ConsensusPathDB tool (http://cpdb.molgen.mpg.de). Heatmaps were generated in R using the heatmap.2 function contained within the gplots package. The Ward method was used for clustering of heatmaps.

#### Raw data accession and panorama public spectral libraries

The mass spectrometric raw data associated with this manuscript may be downloaded from MassiVE at ftp://MSV000087116@massive.ucsd.edu (password: winter). MassIVE ID number: MSV000087116 (password: winter).

### Results

#### Cystine stone development is characterized by a sediment-like precursor, a stone nidus, and growth from the nidus

Mice deficient for either the *Slc3a1* or *Slc7a9* gene develop cystine urinary stones, corresponding to similar stone types in human cystinuria [26]. We utilized these mouse models to study the initiation and progression of these cystinuric stones **(Fig. 1)**. Unlike most urinary human cystine stones, mouse urinary stones manifest primarily in the bladder. This feature of murine stone formation enabled *in vivo* imaging of urinary stone development in an environment without the physical constraints of the renal medulla. The early stages of stone formation are thus clearly discernable by μCT.

**Fig. 1.**
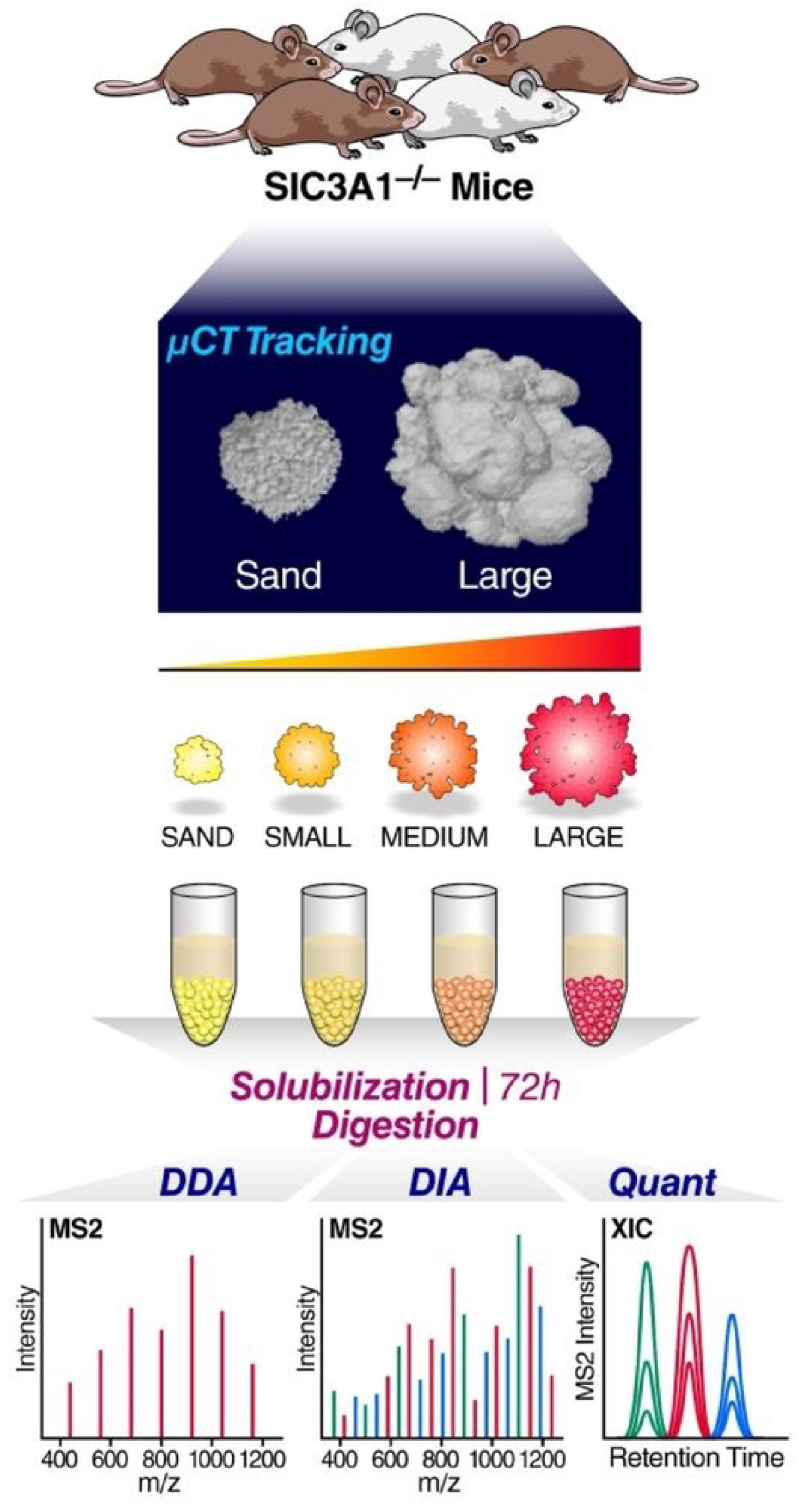
Workflow for tracking and proteomic analysis of *Slc3a1-/-* mouse cystinuric stones. Shows representative images for the tracking of mouse stones using μCT, collecting and solubilizing for mass spectrometry analysis using both DDA and DIA.

Using μCT to examine individual mice, we found that stone formation is initiated by urinary cystine sediment that accumulates in the bladder (**Fig. 2A**). In 15 of 16 mice analyzed, a sediment-like stage was detected prior to the presence of stones. The sediment was characterized by its granular size and shape in Hounsfield units that correspond to the radiodensity of cystine stones. These observations suggest that the sediment is a precursor of stones in mice and that the sediment aggregates in a remodeling process to form distinct stones.

**Fig. 2.**
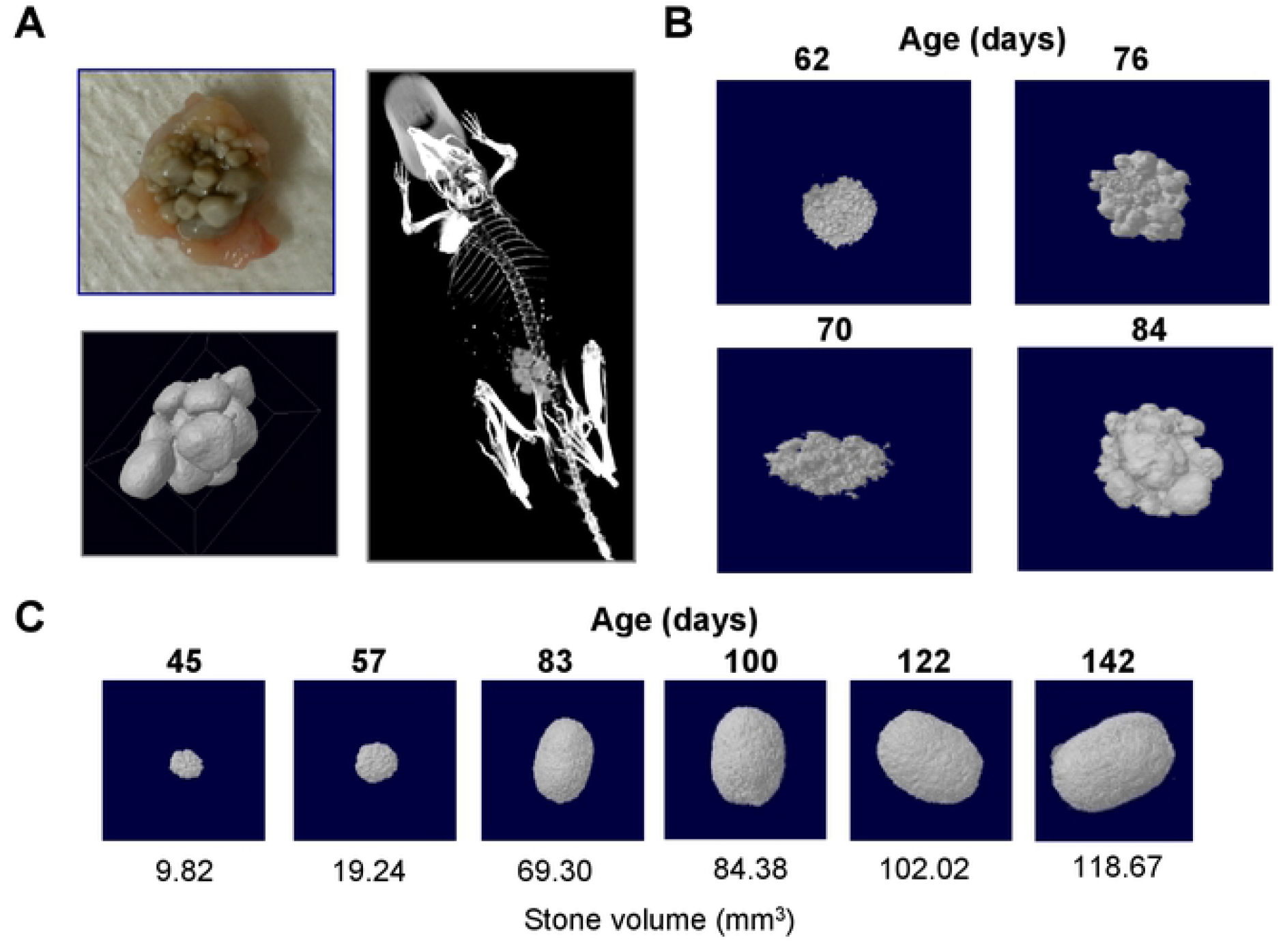
Stages of stone development. (A) μCT imaging and 3D modeling of *in vivo* cystine stone development in a representative *Slc3a1^-/-^* mouse. (B) μCT imaging and 3D modeling of urinary stone growth in an individual *Slc3a1^-/-^* mouse. Stone formation often initiates as sediment in the bladder (P62), and progresses through agglomeration (P70– 76) to form urinary stones (P84); this progression can be seen here. (C) μCT imaging and 3D modeling of a single stone’s growth in mm^3^ over 142 days.

In *Slc3a1^-/-^* mice, stones of different sizes often develop concurrently. We analyzed the growth of individual stones that were discernable by their distinct sizes. We found that cystine stones initiate as individual nidi and accumulate volume in a gradual, appositional process (**Fig. 2B-C**). From here forward, the stones will be referred to as “sand”, “small”, “medium” and “large” stones. From these *in vivo* observations, we propose a model of cystine stone development that depends on accumulation and formation of cystine crystal sediment. A nidus forms from this granular precursor and develops into a developed urinary stone.

#### Cystine stone development is associated with changes in the protein matrix

To understand the contribution of the protein matrix to cystine stone formation, we used mass spectrometry to characterize the proteome of the different stages of stone formation in the cystinuric mouse: the cystine sediment and large cystine stones. To assess the proteins associated with the stone matrix, we used sample preparation protocols for protein extractions from bone [17]. Overall, we identified 1034 unique proteins from all stone types (**S1 Table**). Interestingly, of the 849 proteins in these two stone types, 426 were found in common in sand and large stones (**Fig. 3A**). Using the more accurate relative quantification from our proteomic DIA workflow, we determined significant abundance differences in over 400 proteins across the stone proteome (large vs sand). We found a constantly changing organic matrix in these stones, and interestingly, each size distinction displayed its own signature (**Fig. 3B**). These results suggest that specific urinary proteins and other biomolecules have roles in cystine stone development.

**Fig. 3.**
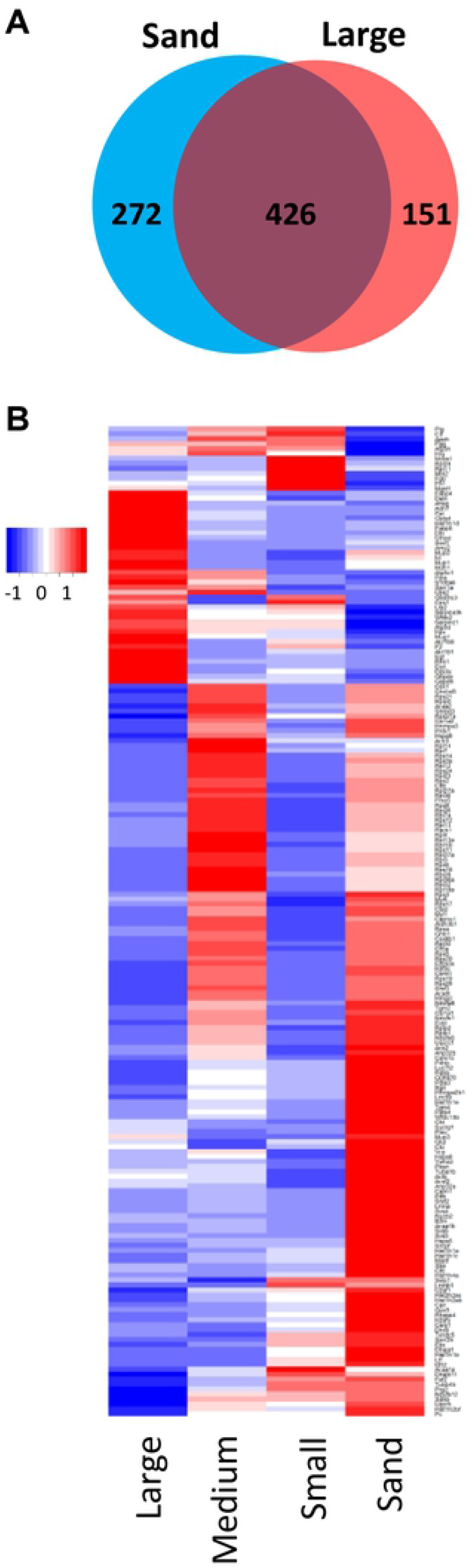
Proteomic analysis of stone development in large and sand kidney stones. (A) Venn diagram comparing the proteins identified in sand-sized stones (698 proteins) versus large stones (577 proteins). We identified 426 proteins in both fractions. (B) Heatmap of all proteins that were significantly changed in abundance in small, medium, and large versus sand stones (q-value < 0.05) by at least 1.5-fold (200 proteins total). Heatmap colors represent the log2 fold-change of each protein versus the median value in the fraction.

We then determined what GO pathways related to the proteins found in the stones were altered (comparing large and sand-type stones). We show both upregulation and downregulation of multiple pathways, including blood coagulation, ribosomal enrichment, metabolism, and RNA processing and transport (**Fig. 4-5**). We also compared the identified proteins from the mouse bladder stones with reported proteomes, such as the human urine proteome (Human Proteome Project, HPP). Of the proteins detected in mouse bladder stones, 643 were described as homologous proteins in human urine (Peptide Atlas human urine HUPO[13, 27–31]) (**S2 Table**). In addition, we compared our list of bladder stone proteins with proteins reported from other bladder or kidney stone studies [27–31]. As in humans, these mice show markers of kidney injury, increased fibrinolysis activity, and dysregulation of proteolysis with the change of both proteases and protease inhibitors. These observations suggest that cystinuric mice share reported proteomic profiles obtained from human urine, bladder and kidney studies, which is highly relevant for the potential future applications of this specific mouse model.

**Fig. 4.**
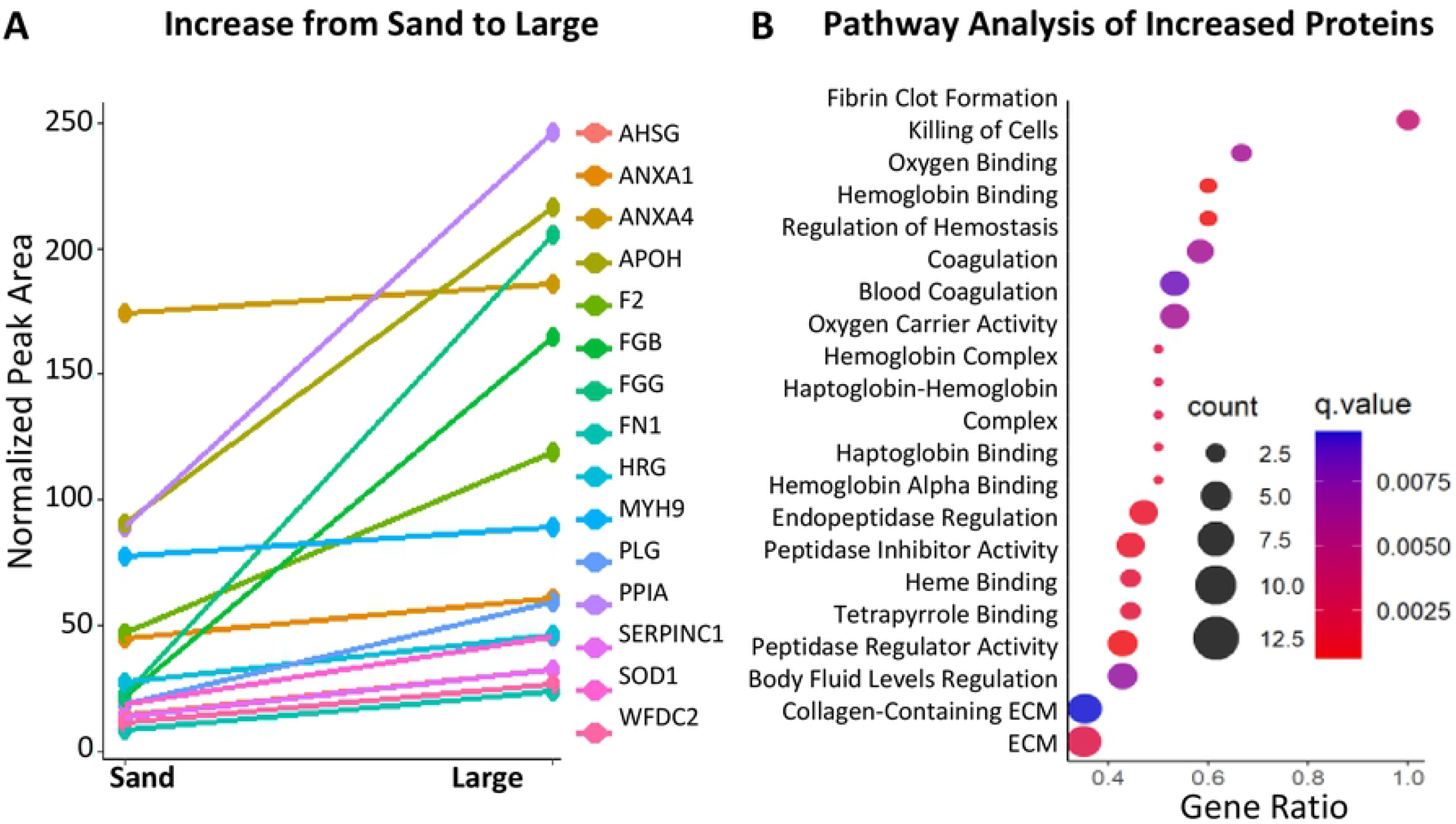
Pathway analysis of significantly changed proteins in sand and large stones. (A) Among upregulated pathways and proteins in large stones, coagulation machinery and protease inhibitors are significantly increased. (B) There is also enrichment for coagulation-related pathways and hemoglobin binding and oxygen binding pathways.

**Fig. 5.**
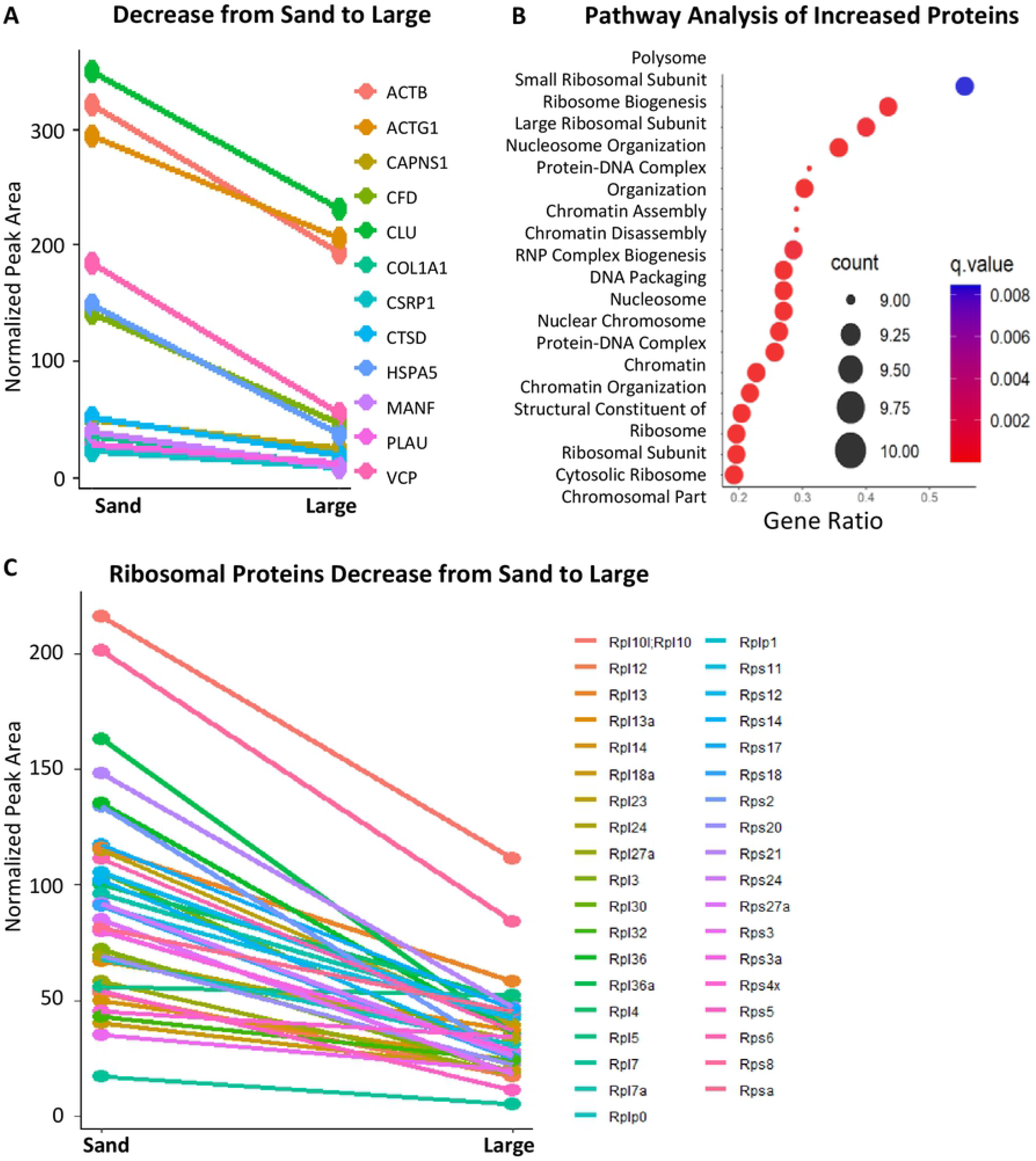
Pathway analysis of significantly changed proteins in sand and large stones. (A) Among downregulated pathways and proteins in large stones, RNA export, ribosomal binding and proteases are significantly decreased. (B) Polysome and ribosomal-related pathways are most significantly enriched, but there is also significant enrichment for chromosomal pathways. (C) Levels of many ribosomal proteins decrease drastically as ‘sand’ continues to grow into the large stones.

#### Early stone formation is associated with significant enrichment of ribosomal proteins

Amounts of ribosomal proteins, such as the ribosomal protein RPL13, were much greater in the sand samples than in the large stone samples (**Fig. 5C**). Initial seeding of a stone may recruit any of the abundant proteins in the urine [11, 32]. A report of all significantly regulated proteins during stone formation and growth can be found in Supplemental Table S1.

#### Significant changes in coagulation factors across stone size progression

Using our understanding of stone formation, we determined how the stones changed as they increased in size. We found that their proteomic signatures changed drastically as they grew from sand to large-sized stones (**Fig. 3**). These distinct sizes allowed us to elucidate where the most drastic proteomic changes occurred. We also found a downregulation in the urokinase responsible for activating plasminogen (PLAU) (**Fig. 5A**). Similarly, the relative abundance of protease proteins was greater in sand than in large stones, shown in the downregulated network (**Fig. 5A**). We identified a small, but statistically significant, upregulation of prothrombin (F2), another protein involved in coagulation, across all stone sizes, compared to sand (**Fig. 6A**). Fibrinogen side chains (e.g., FGB and FGG) and plasminogen (PLG) were significantly more prevalent across all sizes in comparison to sand (**Fig. 6B-D**). These changes in the stone protein profiles possibly reflect and indicate major alterations in the clotting pathway as part of disease progression.

**Fig. 6.**
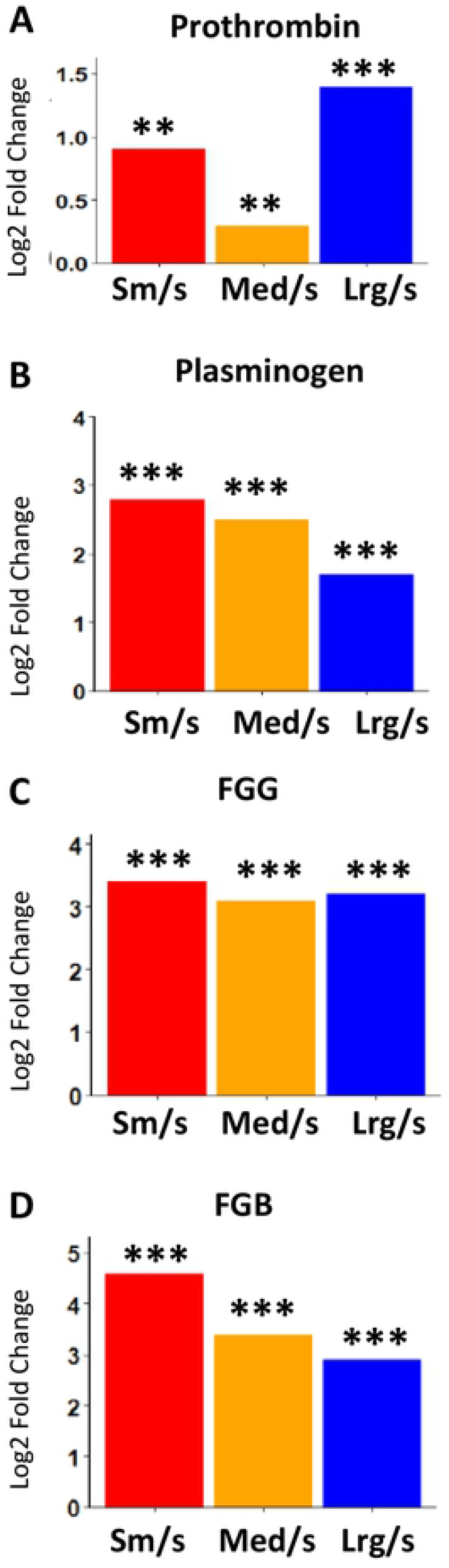
Increases in coagulation machinery detected using log-scale fold-changes. (A) Levels of prothrombin increased across all stone sizes, but most drastically in large stones. (B) Plasminogen shows a large increase in the small stones, but still shows an increase as the stones grow. (C-D) Fibrinogen side-chains (FGG) show large increases across all stone sizes, and beta chains (FGB) show a stepwise decline as the stones increase in size. **=p<0.005; ***=p<0.0005)

#### Protease and protease inhibitor activities are indicative of stone growth

Proteases and protease inhibitors appear most drastically altered in the larger categories. Levels of cysteine protease Calpain 4 and the aspartyl protease Cathepsin D were significantly lower in larger stones than in sand (**Fig. 7A-B**). Alternatively, more protease inhibitor alpha-2-HS-glycoprotein (AHSG) was found in the large stones than in the sand (**Fig. 7C**). Using this trend, we hypothesize that restriction of protease activity may also have a role in stone formation or expansion. Interestingly, SOD1 was increasingly more prevalent as the stones grew (**Fig. 7D**). SOD1 activity is required for processing superoxide, and the effect of its elevated presence in only the largest stones is rather interesting.

**Fig. 7.**
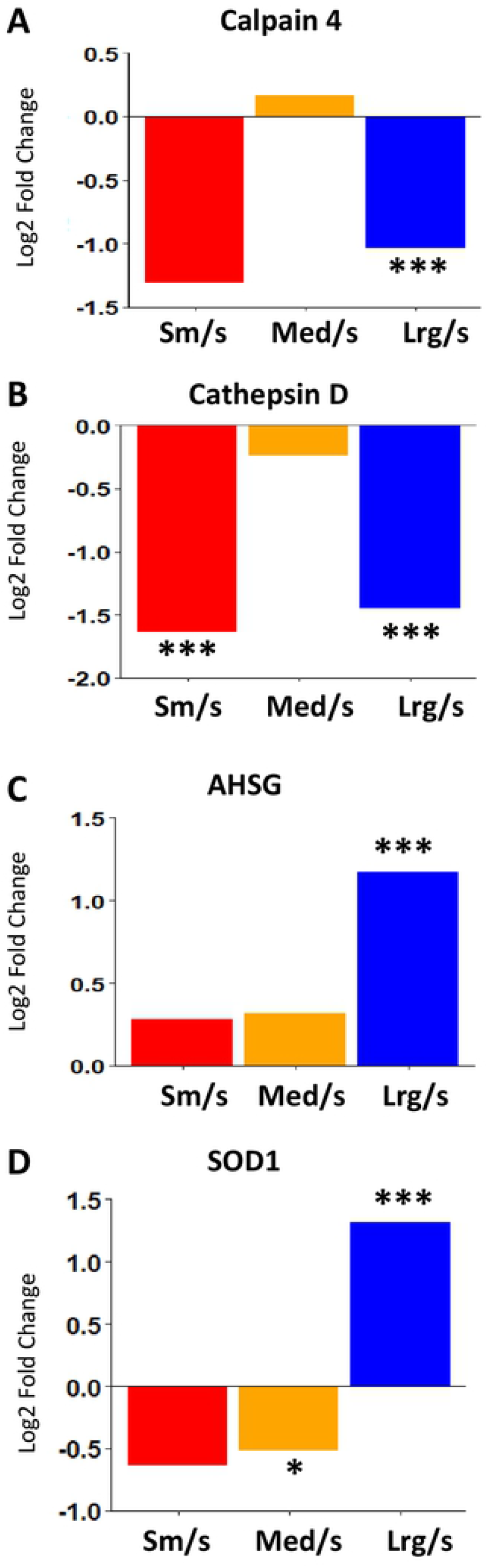
Protein processing and regulatory protein amounts significantly altered in stones when compared to sand. (A-B) Lower levels of proteases Calpain 4 and Cathepsin D are found in large stones than in sand. (C) Protease inhibitor AHSG was increased in large stones. (D) SOD1 follows the same pattern as for AHSG as it is most drastically changed in the large stones, compared to sand. (*=p<0.05; ***=p<0.0005)

### Discussion

To gain insight into the cystine stone development process, we applied longitudinal μCT scanning to characterize early stone formation and initiation in a mouse model of cystinuria (*Slc3a1^-/-^*). These mice lack the transporter required to reabsorb cystine and develop cystinuric stones throughout their life. We then used in-depth proteomic analysis to identify the protein changes associated with the different stages of stone formation and growth. By combining imaging and proteomic analysis of these stages of cystine stone formation, we found stone matrix protein changes that are associated with cystine stone maturation. These formation-associated stone proteins may provide information on stone formation mechanisms and the systems that promote stone development including growth and maturation. We also recapitulated and significantly expanded on the existing data [33, 34], including observations of upregulation of fibrosis and inflammation factors, but we have also captured interesting distinctions in the varying sizes of collected stones that would have not been possible using humans as the primary model. Because current treatments of cystinuria are not effective in the long term, it is necessary to discover new therapeutic targets to treat this disorder.

Many of the bladder stone proteins we identified in this study are also found in human calcium-based stones and in the human urine proteome [11, 32]. Proteins that are overrepresented in the early development of the cystine stones (*sand*) include ribosomal proteins and metabolic factors in the electron transport chain and TCA cycle. In large stones, the relative increased levels of coagulation factors, protease inhibitors, and SOD1 suggest that injury and inflammation may be a normal part of stone formation. Hydrogen peroxide also reacts with free cysteine to form the oxidized dimer cystine, reportedly leading to kidney stone formation [35]. Interestingly, we found SOD1, which generates H2O2 from superoxide, at higher relative abundances in large stones. Accumulation of SOD1 within the stones may promote local conditions that include a higher local H2O2 and lower pH that contributes to stone development. All of these factors need to be investigated further in reference to stone initiation and expansion. However, this is a first study showing how complex and dynamic the protein profiles are inside growing cystine stones. Also, the initial nidus of the urinary stone may cause a level of inflammation that signals for the coagulation response and is bolstered by a lack of relevant proteases and an increase in protease inhibitors to clear constantly amalgamating protein clumps. As a consequence, the stones are never properly dissolved in the body and continue to grow into larger stones. In this study, there were significant increases in protease inhibitors, such as the Serpin family of protease inhibitors, and significant decreases in cysteine proteases, CAPN1 and 4, as well as the aspartyl protease Cathepsin D. Therefore, these proteases cannot clear amalgamating peptides, which could lead to decreased stone solubility in susceptible bladders. Interestingly, patients treated for HIV with Atazanavir have an increased risk of developing stones [36, 37]. Atazanavir is a protease inhibitor that specifically inhibits HIV-1 protease, an aspartyl protease like the Cathepsin proteases. More experiments are needed to identify a method of stone initiation involving these molecules, but these data suggest a link between a lack of protease activity and an increase in stone formation [36, 38]. Our analysis also identified multiple important clotting factors as significantly upregulated in large stones.

The protease inhibitor alpha-2-hs-glycoprotein (AHSG) is also interesting. We found that its presence was increased significantly only in the large stones and was equally found in small and medium, compared to sand. This glycoprotein is upregulated in urine studies linking its expression to an increased risk for kidney disease [39]. AHSG knockdown in mouse models lead to calcification of the vasculature [40]. These mouse models are relatively well studied, but more kidney research is needed to determine any specific role of AHSG in the progression of these disorders. Because the knockout models lead to calcification, AHSG could be involved in a protective molecular mechanism in response to kidney injury. Our data suggest that, as the stones increase in size from medium to large, AHSG accumulates past the levels seen in the other size distinctions. Its role in calcification and known status as a circulating protein make it an important biomarker for multiple kidney- and bladder-related disorders.

Clotting activity is complex and highly regulated through different protein pathways. In this study, we detected significant fold-changes, both increasing and decreasing, across all stone sizes. Some of these factors include components that make up blood clots, fibrinogen side chains, and the machinery regulating their degradation, such as plasmin and antithrombin. Two of the known fibrin subunits (FGG and FGB) were upregulated across all stone sizes. Indeed, patients with chronic kidney disease are more likely to have blood clot clearance issues and show a proteomic signature similar to these stones [33, 34].

We determined that levels of PLG were higher in all stones than in sand stones. Interestingly, PLG showed similar fold-change patterns across stone sizes when compared to fibrin beta chain, which was most drastically increased in the small stones. Increases in the fibrin side-chains tapered off as the stones get larger. PLG, while responsible for degradation of the fibrin subunits, must be cleaved to be activated. This suggests that, even upon an increase in PLG expression, a large amount of fibrin clots could be left intact. This led us to examine the levels of one of the proteins responsible for the cleavage of the inactive plasminogen, urokinase-type plasminogen activator (PLAU). PLAU levels were decreased across all stone sizes, specifically and significantly in the larger stones. However, levels of prothrombin, another crucial component of the clotting machinery, were significantly higher in stones than sand samples. Thrombin is responsible for the formation of the insoluble fibrin clots by mediating the cleavage of fibrinogen to fibrin. Prothrombin as found across all stone sizes, suggesting and reflecting active clotting processes in the disease model. Other stone-associated factors may contribute to the stone matrix proteome.

The significance of these differences between the cystine sediment and sizable formed stones remains to be defined. Our analyses of mouse cystine stone formation reveal that these cystine stones contain a significant organic matrix component, which has not been reported to date. We determined that formation of large stones is preceded by deposition of urinary cystine ‘sand’-like particles, which are likely precursors of the growing stones. By combining imaging analysis and proteomic analysis, we discovered a stone proteome that is constantly changing as stones develop, and we also propose possible mechanisms by which stone formation and growth occur and that could be explored in future studies. These findings could be useful when typical methods of treatment still lead to stone reoccurrence. Here we provide druggable targets in coagulation machinery, protease activity, as well as some metabolic proteins. It is also important to consider stone formation when administering protease inhibitors, such as Atazanavir, as these drugs may impede cystine reabsorption or otherwise contribute to stone formation.

## Acknowledgments

This work was supported by a NCRR shared instrumentation grant 1S10 OD016281 (Buck Institute).

## Author Contributions

Conceptualization: PK, BS, MS, TZ; Data curation: TZ, BS, NaB, JR; Formal analysis: BS, JR, CW, NaB; Funding acquisition: BS, PK; Investigation: JR, TZ, NeB, BS, PK; Methodology: TZ, NeB, BS; Resources: PK, BS; Software: BS, CW, NaB; Supervision: PK, BS; Validation: JR, BS, NaB; Visualization: BS, NaB, CS, JR; Writing – original draft: JR, TZ, NaB, BS; Writing – review & editing: BS, PYD, PK, MS

## Footnotes

The authors declare no competing financial interests.

## Abbreviations

MS1: full scan mass spectrum
MS/MS: tandem mass spectrometry
DTT: dithiothreitol
XIC: extracted ion chromatogram
DDA: data-dependent acquisition
DIA: data-independent acquisition
FDR: false discovery rate
CV: coefficient of variation.

## References

1. Ahmed K, Dasgupta P, Khan MS. Cystine calculi: challenging group of stones. Postgrad Med J. 2006;82(974):799–801. doi: 10.1136/pgmj.2005.044156. PubMed PMID: 17148700; PubMed Central PMCID: PMCPMC2653923.

2. Calonge MJ, Gasparini P, Chillaron J, Chillon M, Gallucci M, Rousaud F, et al. Cystinuria caused by mutations in rBAT, a gene involved in the transport of cystine. Nature genetics. 1994;6(4):420–5. doi: 10.1038/ng0494-420 [doi].

3. Calonge MJ, Nadal M, Calvano S, Testar X, Zelante L, Zorzano A, et al. Assignment of the gene responsible for cystinuria (rBAT) and of markers D2S119 and D2S177 to 2p16 by fluorescence in situ hybridization. Hum Genet. 1995;95(6):633–6. PubMed PMID: 7789946.

4. Mattoo A, Goldfarb DS. Cystinuria. Semin Nephrol. 2008;28(2):181–91. doi: 10.1016/j.semnephrol.2008.01.011. PubMed PMID: 18359399.

5. Ishak R, Abbas O. Penicillamine revisited: historic overview and review of the clinical uses and cutaneous adverse effects. Am J Clin Dermatol. 2013;14(3):223–33. doi: 10.1007/s40257-013-0022-z. PubMed PMID: 23605177.

6. Bagga HS, Chi T, Miller J, Stoller ML. New insights into the pathogenesis of renal calculi. Urol Clin North Am. 2013;40(1):1–12. doi: 10.1016/j.ucl.2012.09.006. PubMed PMID: 23177630; PubMed Central PMCID: PMCPMC4165395.

7. Miller NL, Evan AP, Lingeman JE. Pathogenesis of renal calculi. Urol Clin North Am. 2007;34(3):295–313. doi: 10.1016/j.ucl.2007.05.007. PubMed PMID: 17678981.

8. van Aswegen CH, du Plessis DJ. Pathogenesis of kidney stones. Medical hypotheses. 1991;36(4):368–70. doi: 0306-9877(91)90011-M [pii].

9. Dello Strologo L, Pras E, Pontesilli C, Beccia E, Ricci-Barbini V, de Sanctis L, et al. Comparison between SLC3A1 and SLC7A9 cystinuria patients and carriers: a need for a new classification. J Am Soc Nephrol. 2002;13(10):2547–53. PubMed PMID: 12239244.

10. Santucci L, Bruschi M, Candiano G, Lugani F, Petretto A, Bonanni A, et al. Urine Proteome Biomarkers in Kidney Diseases. I. Limits, Perspectives, and First Focus on Normal Urine. Biomarker insights. 2016;11:41–8. doi: 10.4137/BMI.S26229.

11. Zhao M, Li M, Yang Y, Guo Z, Sun Y, Shao C, et al. A comprehensive analysis and annotation of human normal urinary proteome. Scientific reports. 2017;7(1):3024. doi: 10.1038/s41598-017-03226-6.

12. Kovacevic L, Lu H, Goldfarb DS, Lakshmanan Y, Caruso JA. Urine proteomic analysis in cystinuric children with renal stones. Journal of pediatric urology. 2015;11(4):217.e1–.e2176. doi: 10.1016/j.jpurol.2015.04.020.

13. Bourderioux M, Nguyen-Khoa T, Chhuon C, Jeanson L, Tondelier D, Walczak M, et al. A new workflow for proteomic analysis of urinary exosomes and assessment in cystinuria patients. J Proteome Res. 2015;14(1):567–77. Epub 2014/11/05. doi: 10.1021/pr501003q. PubMed PMID: 25365230.

14. Siddiqui AA, Sultana T, Buchholz NP, Waqar MA, Talati J. Proteins in renal stones and urine of stone formers. Urological research. 1998;26(6):383–8. doi: 10.1007/s002400050073 [doi].

15. Gillet LC, Navarro P, Tate S, Rost H, Selevsek N, Reiter L, et al. Targeted data extraction of the MS/MS spectra generated by data-independent acquisition: a new concept for consistent and accurate proteome analysis. Mol Cell Proteomics. 2012;11(6):O111 016717. Epub 2012/01/21. doi: 10.1074/mcp.O111.016717. PubMed PMID: 22261725; PubMed Central PMCID: PMCPMC3433915.

16. Rardin MJ, Schilling B, Cheng LY, MacLean BX, Sorensen DJ, Sahu AK, et al. MS1 Peptide Ion Intensity Chromatograms in MS2 (SWATH) Data Independent Acquisitions. Improving Post Acquisition Analysis of Proteomic Experiments. Mol Cell Proteomics. 2015;14(9):2405–19. Epub 2015/05/20. doi: 10.1074/mcp.O115.048181. PubMed PMID: 25987414; PubMed Central PMCID: PMCPMC4563724.

17. Jiang X, Ye M, Jiang X, Liu G, Feng S, Cui L, et al. Method development of efficient protein extraction in bone tissue for proteome analysis. J Proteome Res. 2007;6(6):2287–94. doi: 10.1021/pr070056t. PubMed PMID: 17488005.

18. Collins BC, Hunter CL, Liu Y, Schilling B, Rosenberger G, Bader SL, et al. Multi-laboratory assessment of reproducibility, qualitative and quantitative performance of SWATH-mass spectrometry. Nat Commun. 2017;8(1):291. Epub 2017/08/23. doi: 10.1038/s41467-017-00249-5. PubMed PMID: 28827567; PubMed Central PMCID: PMCPMC5566333.

19. Meyer JG, D’Souza AK, Sorensen DJ, Rardin MJ, Wolfe AJ, Gibson BW, et al. Quantification of Lysine Acetylation and Succinylation Stoichiometry in Proteins Using Mass Spectrometric Data-Independent Acquisitions (SWATH). J Am Soc Mass Spectrom. 2016;27(11):1758–71. Epub 2016/09/04. doi: 10.1007/s13361-016-1476-z. PubMed PMID: 27590315; PubMed Central PMCID: PMCPMC5059418.

20. Schilling B, Gibson BW, Hunter CL. Generation of High-Quality SWATH((R)) Acquisition Data for Label-free Quantitative Proteomics Studies Using TripleTOF((R)) Mass Spectrometers. Methods Mol Biol. 2017;1550:223–33. Epub 2017/02/12. doi: 10.1007/978-1-4939-6747-6_16. PubMed PMID: 28188533; PubMed Central PMCID: PMCPMC5669615.

21. Kuhn ML, Zemaitaitis B, Hu LI, Sahu A, Sorensen D, Minasov G, et al. Structural, kinetic and proteomic characterization of acetyl phosphate-dependent bacterial protein acetylation. PLoS One. 2014;9(4):e94816. Epub 2014/04/24. doi: 10.1371/journal.pone.0094816. PubMed PMID: 24756028; PubMed Central PMCID: PMCPMC3995681.

22. Schilling B, Rardin MJ, MacLean BX, Zawadzka AM, Frewen BE, Cusack MP, et al. Platform-independent and label-free quantitation of proteomic data using MS1 extracted ion chromatograms in skyline: application to protein acetylation and phosphorylation. Mol Cell Proteomics. 2012;11(5):202–14. doi: 10.1074/mcp.M112.017707. PubMed PMID: 22454539; PubMed Central PMCID: PMCPMC3418851.

23. Shilov IV, Seymour SL, Patel AA, Loboda A, Tang WH, Keating SP, et al. The Paragon Algorithm, a next generation search engine that uses sequence temperature values and feature probabilities to identify peptides from tandem mass spectra. Mol Cell Proteomics. 2007;6(9):1638–55. Epub 2007/05/30. doi: 10.1074/mcp.T600050-MCP200. PubMed PMID: 17533153.

24. Bruderer R, Bernhardt OM, Gandhi T, Miladinovic SM, Cheng LY, Messner S, et al. Extending the limits of quantitative proteome profiling with data-independent acquisition and application to acetaminophen-treated three-dimensional liver microtissues. Mol Cell Proteomics. 2015;14(5):1400–10. Epub 2015/03/01. doi: 10.1074/mcp.M114.044305. PubMed PMID: 25724911; PubMed Central PMCID: PMCPMC4424408.

25. MacLean B, Tomazela DM, Shulman N, Chambers M, Finney GL, Frewen B, et al. Skyline: an open source document editor for creating and analyzing targeted proteomics experiments. Bioinformatics. 2010;26(7):966–8. Epub 2010/02/12. doi: 10.1093/bioinformatics/btq054. PubMed PMID: 20147306; PubMed Central PMCID: PMCPMC2844992.

26. Font-Llitjos M, Feliubadalo L, Espino M, Cleries R, Manas S, Frey IM, et al. Slc7a9 knockout mouse is a good cystinuria model for antilithiasic pharmacological studies. Am J Physiol Renal Physiol. 2007;293(3):F732–40. doi: 10.1152/ajprenal.00121.2007. PubMed PMID: 17596531.

27. Canales BK, Anderson L, Higgins L, Ensrud-Bowlin K, Roberts KP, Wu B, et al. Proteome of human calcium kidney stones. Urology. 2010;76(4):1017 e13–20. Epub 2010/08/17. doi: 10.1016/j.urology.2010.05.005. PubMed PMID: 20709378.

28. Canales BK, Anderson L, Higgins L, Frethem C, Ressler A, Kim IW, et al. Proteomic analysis of a matrix stone: a case report. Urol Res. 2009;37(6):323–9. Epub 2009/09/05. doi: 10.1007/s00240-009-0213-5. PubMed PMID: 19730843.

29. Okumura N, Tsujihata M, Momohara C, Yoshioka I, Suto K, Nonomura N, et al. Diversity in protein profiles of individual calcium oxalate kidney stones. PLoS One. 2013;8(7):e68624. Epub 2013/07/23. doi: 10.1371/journal.pone.0068624. PubMed PMID: 23874695; PubMed Central PMCID: PMCPMC3706363.

30. Jou YC, Fang CY, Chen SY, Chen FH, Cheng MC, Shen CH, et al. Proteomic study of renal uric acid stone. Urology. 2012;80(2):260–6. Epub 2012/04/21. doi: 10.1016/j.urology.2012.02.019. PubMed PMID: 22516363.

31. Liu JD, Liu JJ, Yuan JH, Tao GH, Wu DS, Yang XF, et al. Proteome of melamine urinary bladder stones and implication for stone formation. Toxicol Lett. 2012;212(3):307–14. Epub 2012/06/13. doi: 10.1016/j.toxlet.2012.05.017. PubMed PMID: 22688180.

32. Kolbach-Mandel AM, Mandel NS, Hoffmann BR, Kleinman JG, Wesson JA. Stone former urine proteome demonstrates a cationic shift in protein distribution compared to normal. Urolithiasis. 2017;45(4):337–46. Epub 2017/03/21. doi: 10.1007/s00240-017-0969-y. PubMed PMID: 28314883; PubMed Central PMCID: PMCPMC5511579.

33. Huang M-J, Wei R-B, Wang Y, Su T-Y, Di P, Li Q-P, et al. Blood coagulation system in patients with chronic kidney disease: a prospective observational study. BMJ open. 2017;7(5):e014294. doi: 10.1136/bmjopen-2016-014294.

34. Leslie SW, Nazzal L. Renal Calculi (Cystinuria, Cystine Stones). StatPearls. Treasure Island (FL): StatPearls Publishing; 2017.

35. Trivedi P, Kumar RK, Iyer A, Boswell S, Gerarduzzi C, Dadhania VP, et al. Targeting Phospholipase D4 Attenuates Kidney Fibrosis. J Am Soc Nephrol. 2017;28(12):3579–89. Epub 2017/08/18. doi: 10.1681/ASN.2016111222. PubMed PMID: 28814511; PubMed Central PMCID: PMCPMC5698063.

36. Gentle DL, Stoller ML, Jarrett TW, Ward JF, Geib KS, Wood AF. Protease inhibitor-induced urolithiasis. Urology. 1997;50(4):508–11. Epub 1997/10/24. doi: 10.1016/S0090-4295(97)00401-9. PubMed PMID: 9338723.

37. Sundaram C, Saltzman B. Urolithiasis Associated with Protease Inhibitors. Journal of endourology / Endourological Society. 1999;13:309–12. doi: 10.1089/end.1999.13.309.

38. Schwartz BF, Schenkman N, Armenakas NA, Stoller ML. Imaging characteristics of indinavir calculi. J Urol. 1999;161(4):1085–7. Epub 1999/03/19. PubMed PMID: 10081843.

39. Piazzon N, Bernet F, Guihard L, Leonhard WN, Urfer S, Firsov D, et al. Urine Fetuin-A is a biomarker of autosomal dominant polycystic kidney disease progression. Journal of Translational Medicine. 2015;13(1):103. doi: 10.1186/s12967-015-0463-7.

40. Westenfeld R, Schäfer C, Smeets R, Brandenburg VM, Floege J, Ketteler M, et al. Fetuin-A (AHSG) prevents extraosseous calcification induced by uraemia and phosphate challenge in mice. Nephrology Dialysis Transplantation. 2007;22(6):1537–46. doi: 10.1093/ndt/gfm094.

